# Deep Learning identifies new morphological patterns of Homologous Recombination Deficiency in luminal breast cancers from whole slide images

**DOI:** 10.1101/2021.09.10.459734

**Authors:** Tristan Lazard, Guillaume Bataillon, Peter Naylor, Tatiana Popova, François-Clément Bidard, Dominique Stoppa-Lyonnet, Marc-Henri Stern, Etienne Decencière, Thomas Walter, Anne Vincent Salomon

## Abstract

Homologous Recombination DNA-repair deficiency (HRD) is a well-recognized marker of platinum-salt and PARP inhibitor chemotherapies in ovarian and breast cancers (BC). Causing high genomic instability, HRD is currently determined by *BRCA1/2* sequencing or by genomic signatures, but its morphological manifestation is not well understood. Deep Learning (DL) is a powerful machine learning technique that has been recently shown to be capable of predicting genomic signatures from stained tissue slides. However, DL is known to be sensitive to dataset biases and lacks interpretability. Here, we present and evaluate a strategy to control for biases in retrospective cohorts. We train a deep-learning model to predict the HRD in a controlled cohort with unprecedented accuracy (AUC: 0.86) and we develop a new visualization technique that allows for automatic extraction of new morphological features related to HRD. We analyze in detail the extracted morphological patterns that open new hypotheses on the phenotypic impact of HRD.

---

The advent of Deep Learning has revolutionized biomedical image analysis and in particular digital pathology. Traditionally, the majority of methods developed in this field were dedicated to computer-aided diagnosis, where the objective is to partially automatize human interpretation of slides, in order to help pathologists in their diagnosis task, e.g. the detection of mitoses (1), or the identification of metastatic axillary lymph nodes (2,3). Beyond the automatization of manual inspection, Deep Learning has also been successfully applied to predict patient variables, such as outcome (4), and to predict molecular features, such as gene mutations (5,6), expression levels (7) or genetic signatures (5,8). Despite these results of unprecedented quality, one of the major drawbacks of Deep Learning algorithms is their black-box character: because the features are automatically extracted, it is difficult to know how a decision was made. This has two major consequences: first, it is difficult to identify potential confounders, i.e. variables that correlate with the output due to the composition of the data set and that are predicted instead of the intended output variable. Second, even in the absence of statistical artifacts, understanding how the decision was generated in the first place can point to interesting mechanistic hypotheses and to patterns in the image that have so far been overlooked.

One way to overcome the latter problem is to use hand-crafted biologically meaningful features (8). This however requires an extraordinary effort in terms of annotation. Here, we take a conceptually different approach. Instead of working in a pan-cancer setting on a large number of signatures, we concentrate on one single medically highly relevant signature in one cancer type on a controlled data set, where we can correct for potential biases. In order to understand how the Deep Learning decision is generated and which morphological patterns are related to the output variable, we propose a novel visualization and interpretation technique that paves the way to “machine teaching”, i.e. a data driven approach to identify phenotypic patterns related to genomic signatures, thus pointing to new mechanistic hypotheses.

In order to demonstrate the power of this strategy, we focus on predicting Homologous Recombination Deficiency (HRD) in Breast Cancer (BC). Worldwide, 2.1 million women are newly diagnosed per year with BC which is a leading cause of cancer-related death. Improvement of metastatic breast cancer treatment is therefore of highest priority. BC is a heterogeneous disease with four major molecular classes (luminal A, B, HER2 enriched, and triple-negative breast cancer [TNBC]) benefiting from different therapeutic approaches. If early BC patients have an overall survival of 70 to 80%, metastatic disease is incurable with a short duration of survival (9). Homologous Recombination (HR) is a major and high-fidelity repair pathway of DNA double-strand breaks. Its deficiency, HRD, results in high genomic instability (10) and occurs through diverse mechanisms, including germline or acquired mutations in DNA repair genes, most frequently *BRCA1, BRCA2* or *PALB2*, or through epigenetic alterations of *BRCA1* or *RAD51C*. Importantly, HRD leads to a high sensitivity to polyADP-ribose polymerase inhibitors (PARPi) *in vitro* (11,12). PARPi have been shown to improve metastatic breast cancer progression free survival (13,14). Several methods have been developed to detect HRD, including genomic instability profiling, mutational signatures, or integrating structural and mutational signatures (15–19). However, HRD is currently diagnosed in clinical practice by DNA repair genes sequencing and genomic instability patterns (genomic scar) such as the LST signature (15) or the HRD MyChoice® CDx test (Myriad Genetics). *BRCA1* and *BRCA2* mutations are known predictive markers for response to PARPi (10,14) and platinum salt (20) and the somatic HRD has been more recently recognized as a predictive marker for PARPi in ovarian (10) and breast cancer (21). But neither a specific routinely assessed phenotype nor a morphological pattern indicates the presence of HRD. The majority of hereditary *BRCA1* cancers are TNBC and up to 60-69% of sporadic TNBC harbor a genomic profile of HRD (15,22,23). However, the majority of hereditary *BRCA2* cancers are luminal (24) and HRD also exists in sporadic luminal B (21,25) or in HER2 tumors (26,27). Of note, germline or sporadic alterations of *BRCA* harbor indistinguishable genomic alterations in triple-negative or in luminal tumors (28,29). In that context, a reliable and accurate test is mandatory to select patients for PARPi and platinum salt treatments. Whereas the screening for germline or somatic *BRCA1* and *BRCA2* mutations is feasible for all TNBC (18% of all BC cases), it represents a real challenge in clinical practice if extended to all luminal B tumors (35% of all BC). The recent results of the Olympia trial emphasize the need for an efficient method of screening for BRCA1 and 2 mutations across all breast cancer phenotypes(14). This strategy moreover does not identify the whole diversity of genetic causes of HRD.

In this study, we present an image-based approach to predict HR status from Whole Slide Images (WSI) stained with Hematoxylin Eosin (HE) using deep learning, from a large retrospective series of luminal and triple-negative breast carcinomas with a genomically defined HR status, from a single cancer center. In particular, we show that careful correction for potential biases is essential for such studies and demonstrate the relevance of a correction strategy. Finally, we develop a novel interpretation algorithm that allows the visualization of decisive previously unknown patterns and leads to new hypotheses on disease-relevant genotype-phenotype relationships.

## Results

### A Deep Learning architecture to predict HRD from Whole Slide Images (WSI)

We scanned the most representative HE stained tissue section of the surgical resections specimens of breast cancer from 715 patients with known HR status. The series was composed of 309 Homologous Recombination Proficient (HRP) tumors and 406 Homologous Recombination Deficient tumors (Supplementary Table 1).

Due to their enormous size, analysis of WSI typically relies on the Multiple Instance Learning (MIL) paradigm (30–33). MIL techniques only require slide-level annotations and share the overall architecture (see Figure 1) consisting of 4 main steps: tiling and encoding, tile-scoring, aggregation, and decision.

**Figure 1:**
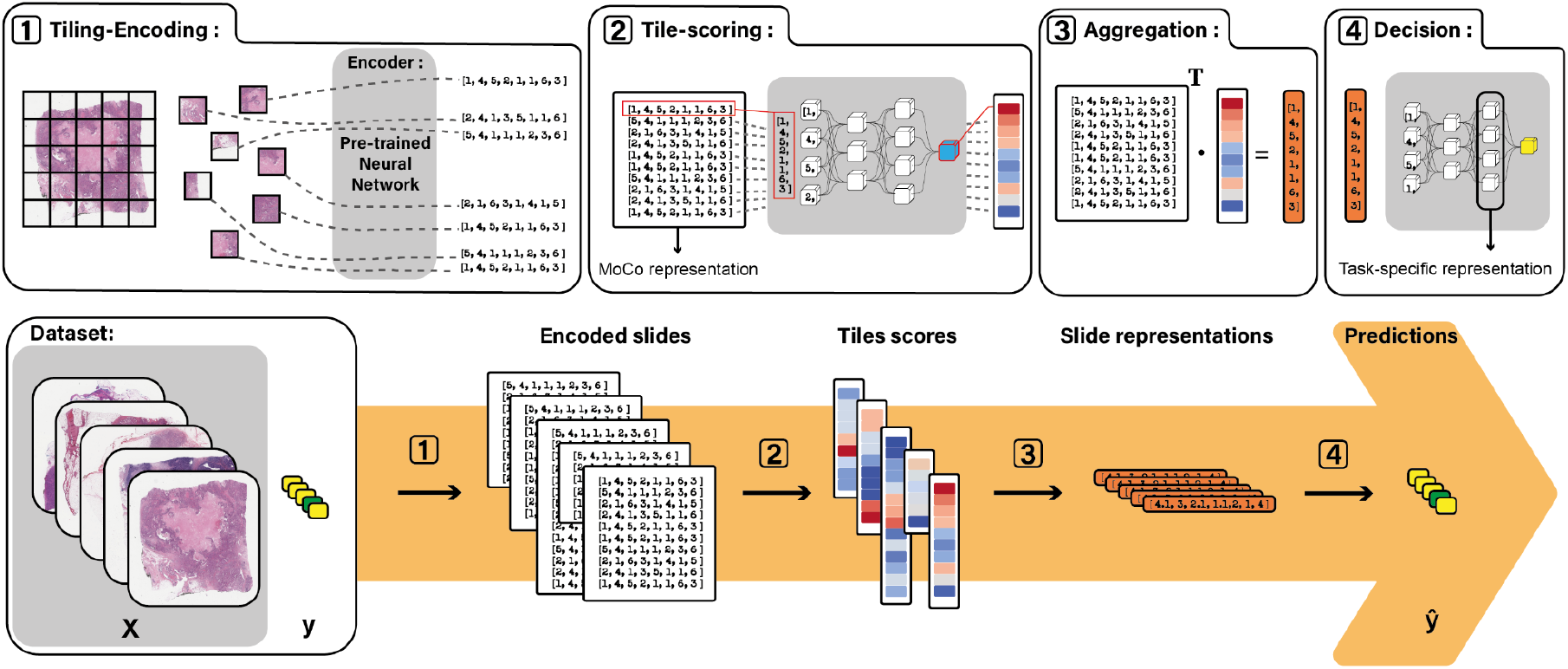
From WSI to prediction. Four major components are used in this end-to-end pipeline. First, the WSI (*X*) are tiled, the tissue parts are automatically selected, and the resulting tiles are embedded into a low-dimensional space (block 1). The embedded tiles are then scored through the attention module (2). An aggregation module outputs the slide level vector representative (3) that is finally fed to a decision module (4) that outputs the final prediction. When training, the binary cross-entropy loss between the ground truth *y* and the prediction is *ŷ* computed and back-propagated to update the parameters of the modules. Both the decision module and the attention module are multi-layer perceptrons, the encoder is a ResNet18 and the aggregation module consists of a weighted sum of the tiles, the weights being the attention scores.

The WSI was divided into tile images (dimension: 224×224 pixels) arranged in a grid. Background tiles are removed, tissue tiles are encoded into a feature vector. Instead of using representations trained on natural image databases and unlike most studies in this domain, we used the self-supervised technique Momentum Contrast (MoCo (34); see Methods). This method consists in training a Neural Network to recognize images after transformations, such as geometric transformations, noise addition and color changes. By choosing the kind and strength of transformations, we can impose invariance classes, i.e. variations in the input that do not result in different representations. The feature vector of each tile was then mapped to a score by a neural network. The slide representation was obtained by the sum of the individual tile representations, weighted by the learned attention scores (30). Finally, the slide representation was classified by the decision module (see Figure 1). We optimized hyperparameters by a systematic random search strategy (see Methods). For hyperparameter setting and performance estimation, we used nested 5-fold cross-validation, which allows us to obtain realistic performance estimations. All reported performance results are averaged over 5 independent test folds (see Methods).

### HRD prediction with correction for potential biases

We applied this method to predict HRD from the WSI in the TCGA cohort, and obtained results (AUC=0.71; see Figure 2.c) in line with previous reports (8,35,36). While the TCGA is an invaluable resource for pan-cancer studies in genomics and histopathology, it is often seen rather as a starting point whose results need to be corroborated by other cohorts (37). Furthermore, the TCGA contains images from many centers around the world with potentially different sample preparation and image acquisition protocols. While this technical variability might reflect to some degree what could be expected in clinical practice for multiple institutions, we hypothesized that to prove the predictability of HRD independently from potential technical and biological biases, as well as an in-depth study of morphological patterns related to HRD, it might be advantageous to work on a more homogeneous data set, where we can carefully control for potential technical and biological confounders. We thus turned to our in-house dataset, the Curie dataset (see Methods), with data from 715 patients.

**Figure 2:**
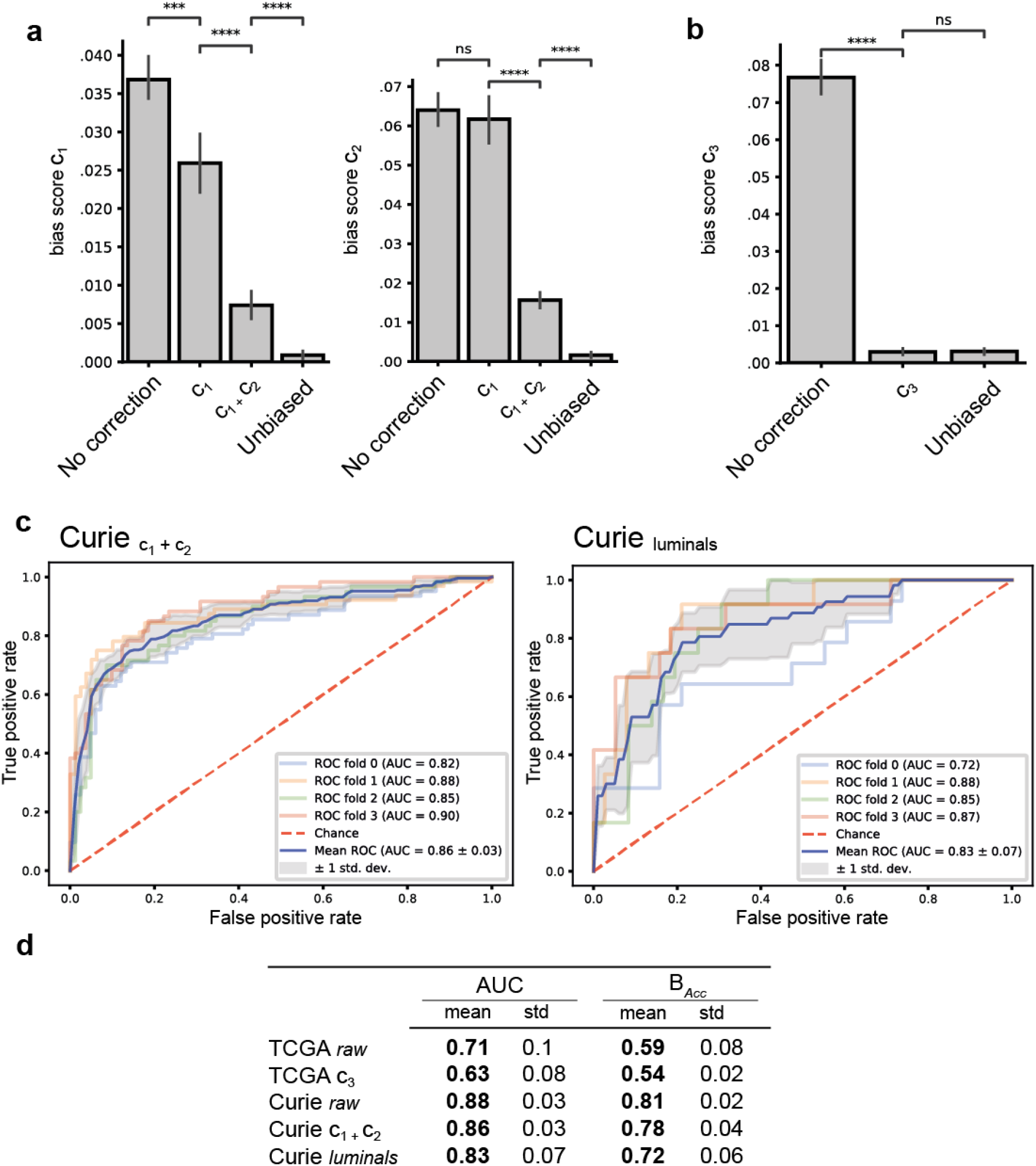
Bias corrections and prediction performances. **a-b**, estimation of the bias score of two technical confounder (*c*_1_, *c*_2_) and one biological confounder (*c*_3_) for the Curie Dataset (**a**) and the bias score of the confounder *c*_3_ for the TCGA dataset (**b**) for different correction strategies. A Mann-Whitney-Wilcoxon test two-sided with Bonferroni correction is performed for each pair of correction strategies. ns: non-significant p>0.05, *: p<0.05, **: p<0.01, ***: p<1e-3, ****: p<1e-4 **c-d**, performance results. Name of each model indicates the origin of its training set. Indices indicate the correction applied through strategic sampling (Curie_*c*1_ has been debiased with respect to *c*_1_). Curie_luminals_ corresponds to the model trained on a subset containing only luminal tumors. **c**: ROC curve of the models trained on the Curie dataset correcting for technical bias (Curie_*c*1_) or for technical biases and *c*_3_ (Curie_luminals_). **d**: summary tables of performance metrics. AUC : Area Under The (receiver operating characteristics) Curve, *B*_Acc_: balanced accuracy.

Training a Neural Network (NN) to predict HRD on the raw data set, we observed an important increase in prediction performances, as compared to the TCGA (AUC=0,88; see Figure 2.c). As the cohort was generated over 25 years, two experimental variables (*c*_1_, *c*_2_, see Methods) representing slight changes in experimental protocols have been identified as potential confounders. To measure the confounding effects of these variables on the model predictions, we developed a bias score (see methods). This score is close to 0 in the unbiased case. We found that model predictions were indeed biased by these two confounders (see Figure 2.a).

We then devised a sampling strategy that mitigates biasing during training. Bias mitigation is an increasingly important line of research in machine learning. For instance, it is a well-known problem in training predictive models for functional magnetic resonance imaging (fMRI) data, where the age of the patient has been shown to be an important confounder (38). While several techniques for bias mitigation exist (39–42), a recent comparison (43) indicates that strategic sampling is the method of choice if the distribution is not too imbalanced. Strategic sampling aims at ensuring that irrespectively of the composition of the training set, each batch presented to the neural network is composed of roughly the same number of samples for each value combination of output and confounding variable. Correcting for *c*_1_ and *c*_2_ resulted in a 4-fold reduction of the bias-score in comparison to the uncorrected model and a slightly lower accuracy (AUC=0.86, Figure 2.c).

In addition to these technical confounders, we identified the molecular subtype of the tumor to be a potential biological confounder. Successful correction of this biological confounder in the TCGA (see figure 2.b) led however to a dramatic drop in performance (AUC=0.63). This result suggests that NN trained on the TCGA for HRD prediction without stratification or bias correction, might actually predict to some extent the molecular subtype, which is also a predictable variable (AUC=0.89). This shows that the molecular subtype is indeed a biological confounder. In our in-house data set, we decided to build a subtype-specific NN that specifically predicts HRD for luminal BC, instead of applying bias mitigation. The reason for this decision was three-fold: first, we argued that a dataset focusing on only one molecular subtype was more likely to reveal the underlying patterns exclusively related to HRD; second, HRD prediction in luminal BC is of particular importance for clinical practice, as very few morphological patterns are known to be related to HRD in luminal BC, the most frequent breast cancer phenotype, and third, the relatively low number of TNBC in our data set made strategic sampling on three confounding variables challenging. Therefore, we composed a dataset containing only luminal BC and setting both technical confounders, leading us to keep 259 BC WSI (191 HRD tumors, 66 HRP tumors). We obtained a good, albeit slightly lower performance of this bias-corrected NN (AUC=0.83, Figure 2.c; Figure 2.d). The trained model carefully freed from both technical and biological biases was then used for the identification of morphological patterns described in the next section.

### Visualization reveals HRD specific tissue patterns

In order to understand which phenotypic patterns are related to HRD on the WSI, we turned to visualization techniques for NN. The used MIL framework is equipped with an inherent visualization mechanism: the second module of the algorithm, the tile-scoring module, is in fact an attention module that assigns to each tile an attention score that determines how much a given tile will contribute to the slide representation (and thus to the decision). Attention scores are often used for visualization in the field of pathology (3,44–46), either in the form of heatmaps in order to localize the origin of the relevant signals or in the form of galleries of tiles of interest (tiles with highest attention scores).

However, attention scores do not per se extract the tiles that are related to a certain output variable; they just reflect that the tile is to be taken into consideration in the decision. In particular in the case of genetic signatures, where we would expect that the output variables can be related to several morphological patterns, analyzing only the attention scores might be limited. In the case of HRD prediction, this intuition is corroborated by the results presented in Figure 3: selecting the tiles with the highest attention scores does not allow to identify tiles related to HRD or HRP, respectively. Moreover, a low-dimensional representation, obtained by Uniform Manifold Approximation and Projection (UMAP) (47) of the tile descriptors does not show any grouping that seems to relate to the output variable.

**Figure 3 :**
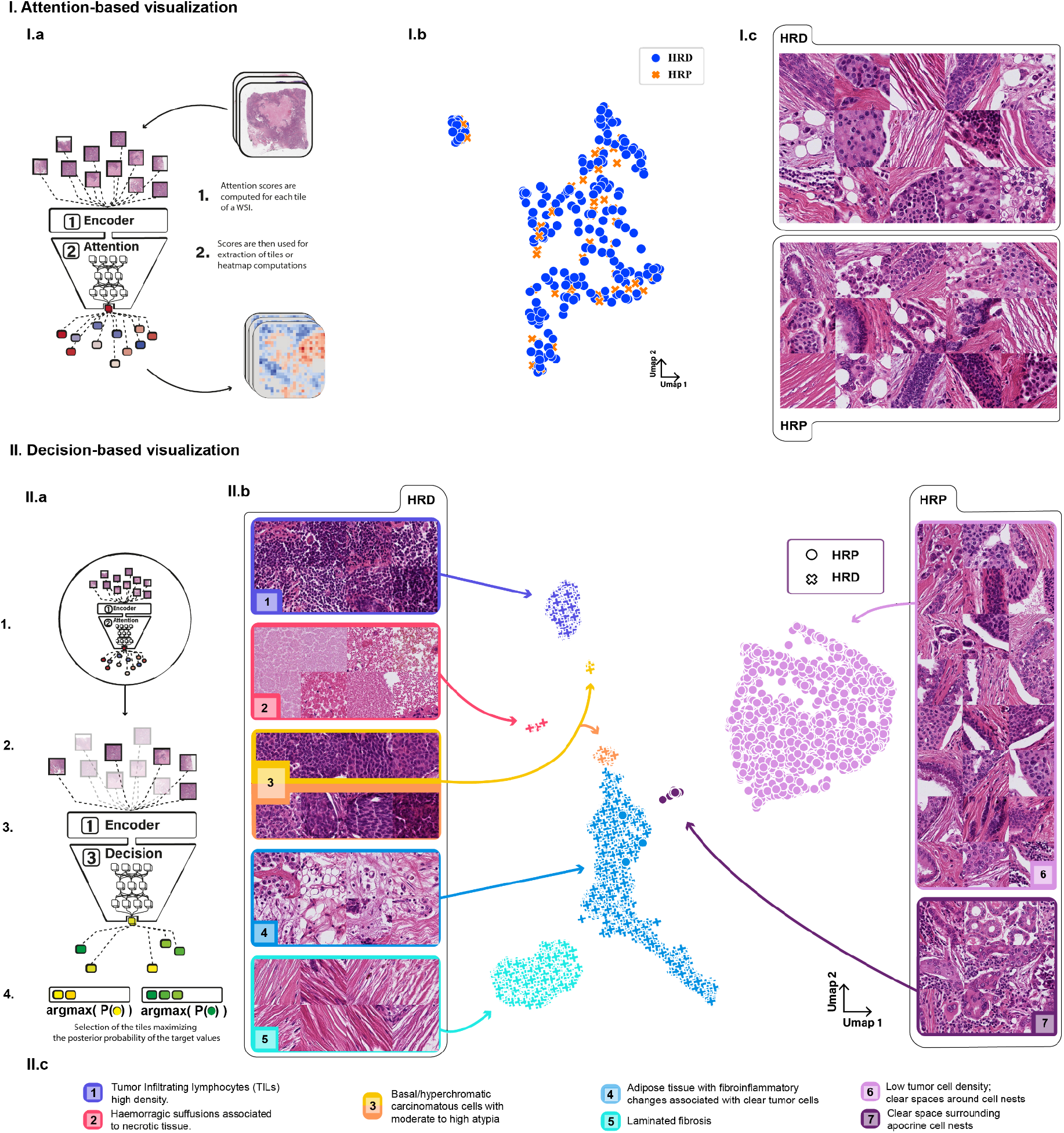
I. Attention-based visualization. does not discriminate between HRD and HRP. **I. a**, Mechanism of the attention-based visualization. The attention score of a tile is used as a direct proxy of its importance in the prediction of the WSI. **I.b**, UMAP projection of the highest attention ranked tiles of the Curie WSIs classified as HRP (orange crosses) and HRD (blue circles). **I.c** Randomly sampled tiles among the HRP and HRD tiles. The tiles are located in the tumor, however, neither clear clusters nor visual differences are present between HRD and HRP tiles. **II. Decision-based visualization. II.a**, Mechanism of the decision-based visualization. 1, Each tile in the whole dataset is scored by the attention module. 2, The best scoring tiles are selected as candidate tiles. 3, The selected tiles are presented to the decision module, the probability of each of these tiles being HRD or HRP (yellow or green) is kept. 4, Finally, the K tiles with maximal probability for either HRD / HRP are selected. **II.b**, Morphological map of the HR status. Each dot is the UMAP projection of a tile extracted by the decision-based visualization method. Crosses are tiles with high probability of being HRD; circles are tiles maximizing HRP prediction. Each cluster has been linked to a morphological phenotype by two expert pathologists. We identified six different morphological phenotypes associated with the HRD and two associated with the HRP. The exhibited tiles have been randomly sampled among each cluster. **II.c** Pathological interpretation of the clusters presented in II.b.

Given these limitations, we propose a new visualization protocol that allows us to extract the tiles that are directly associated with a particular slide-level label. As the slide representation is the weighted sum of the tile representations, we applied the decision module, specifically trained to classify slide representations between HRD and HRP, to the individual tile representations. This gives us a score for each tile that can be interpreted as the (tile) probability of being HRD or HRP (see Methods for details). Selecting the tiles with the highest posterior probability for HRD and HRP respectively, and projecting the tile representations of this selection to a low dimensional space leads to the emergence of distinct clusters corresponding to different tumor tissue patterns with a clear relation to HRD or HRP and therefore providing a morphological map of HRD (Figure 4).

**Figure 4:**
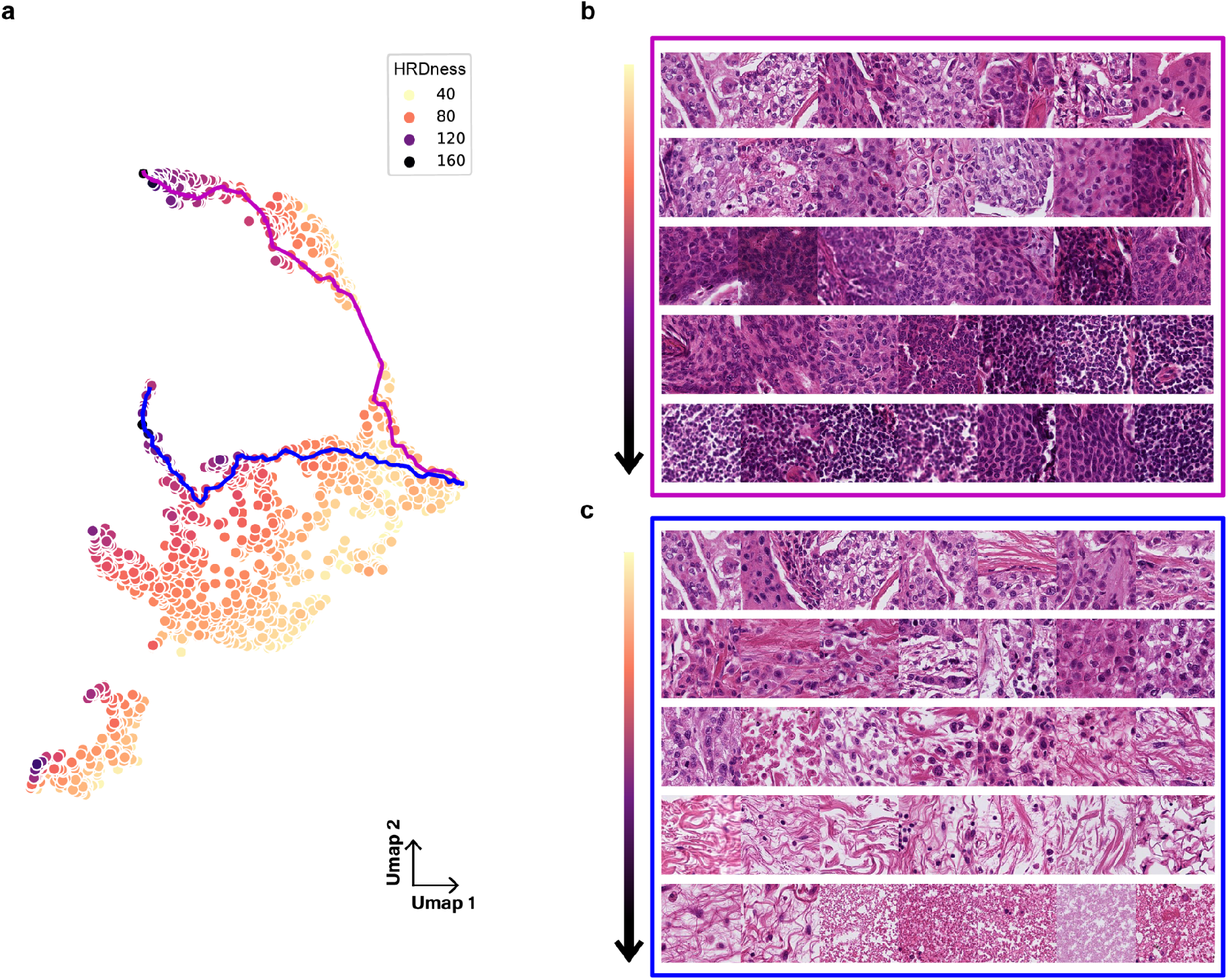
Illustration of 2 Phenotypic HRD-ness trajectories. **a:** UMAP projection of the HR status specific representation of the meaningful tiles relative to the HRD. HRD-ness is the score given to each tile by the HRD output neuron. Two tile trajectories have been extracted (blue and magenta) starting from the same low HRD-ness region, each leading to a different high HRD-ness region. **b, c:** Tiles sampled along each of the trajectories. They are ordered from low HRD-ness to high HRD-ness and read from left to right and from up to bottom. **b**. Magenta trajectory, toward densely cellular tumors or inflammatory cells. **c:** Blue trajectory, toward fibroinflammatory tumor changes and hemorrhagic suffusions.

Two expert pathologists labeled these clusters. The HRD signal relied on several clusters: HRD tumors present a high tumor cell density, with a high nucleus/cytoplasm ratio and conspicuous nucleoli. They also show regions of hemorrhagic suffusion associated with necrotic tissue. In the stroma, the HRD signal revealed the presence of striking laminated fibrosis and as expected relied on high Tumor-Infiltrating Lymphocytes (TILs) content. Lastly, one large cluster contained a continuum of several phenotypes, namely adipose tissue intermingled with scattered and clear tumor cells, histiocytes, and plasma cells. In contrast, the HRP signal was mostly carried by one cluster characterized by low tumor cell density, the cells being moderately atypical and tumor cell nests separated from the stroma by clear spaces. Notably, it included a few invasive lobular carcinomas (all the tiles per cluster are available in the supplementary figures 1-11).

Our approach thus suggests that these tissue phenotypes are hallmarks of HRD. In contrast to triple-negative tumors that have been described as high-grade, rich in TILs, with pushing margins (48), no specific pathological patterns of luminal breast g*BRCA 1 or 2* cancers were identified except a high grade with a frequent absence of tubules formation and pushing margins. However, these features did not allow a robust identification of HRD luminal tumors in clinical practice (49–51). Here, thanks to our visualization tool and our unbiased deep-learning analysis, we identified new features linked to HRD in luminal breast cancers. In order to test this analysis result, the TIL density and nuclear grade were evaluated for each tumor of the in-house dataset by an expert pathologist. As predicted by our algorithm, TILs and nuclear grade were positively associated with the HR status of the tumor in the luminal subset (mean TILs HRD: 29, mean TILs HRP: 17, t-test-pvalue : 0.017; mean nuclear grade HRD: 2.7, mean nuclear grade HRP: 2.3, Xi2-pvalue : 1.2 e-6).

Our NN works with different internal representations. While the tile representations provided by MoCo permit the emergence of phenotypic similarity clusters (Figure 4), internal representations closer to the decision module encode information relevant for HRD. The representation in the penultimate layer can therefore be interpreted as encoding “HRD-ness” of the tiles. Figure 4 illustrates a low dimensional representation of this HRD-ness for the same tiles as those present in Figure 4, where point color represents the HRD-score (tile probability to be classified as HRD). From there, we have extracted two tile trajectories, going from low HRD-ness to high HRD-ness. The magenta trajectory illustrates the successive visual changes corresponding to an increase in tumor cells or inflammatory cells density (from low-density tiles to high-density tiles with large nuclei, nuclear atypia and infiltrative lymphocytes). The blue trajectory shows conversely a decrease in tumor cells density replaced successively by an inflammatory reaction and apoptotic cells, loose fibrosis and hemorrhagic suffusion associated with necrosis. These different trajectories illustrate the manifestations of HRD and show the pleiotropic character of the induced phenotypes. Moreover, the highlighted gradation of these phenotypes opens the path to a possible reading grid of WSIs for pathologists.

## Discussion

In this study, we set out to predict the HR status in breast cancer from H&E stained WSI and to analyze the phenotypic patterns related to HRD. The prediction of HRD is an important challenge in clinical practice. The use of PARP inhibitors for breast cancer patients was initiated for metastatic TNBC patients with germline mutations of *BRCA1* or *BRCA2*. However, *BRCA2*, as well as *PALB2* and a minority of *BRCA1* cancer patients, develop luminal tumors. The necessity of predicting HRD is therefore not limited to TNBC, but extends also to luminal BC. On the other hand, luminal BCs represent a far more frequent group than TNBC. For this reason, systematic screening of HR gene alterations for luminal cancers will be problematic and in many countries even infeasible due to both economic and logistic issues. Therefore, preselection of patients with a high probability of being HR deficient by analysis of WSI is a cost-efficient strategy that has so far only been hampered by the lack of knowledge about HRD specific morphological patterns in luminals. Indeed, only high grades and to a lower extent pushing margins have previously been reported to be associated with HRD. In this context, the identification of HRD from WSI by deep learning and the identification of related morphological patterns could both facilitate the preselection of breast cancers for molecular determination of HRD, which is particularly important for luminal cancers.

The TCGA provides a precious data set to train models for the prediction of genetic signatures from H&E data (5,8). While we obtained promising results for the prediction of HRD on the TCGA dataset in line with previous reports, we found that this result was partly due to the fact that the molecular subtype acts as a biological confounder. This was particularly problematic as we wanted to investigate the morphological signature of HRD. Of note, the existence of biological and technical confounders is presumably not limited to HRD prediction, but may concern many genetic signatures. The use of carefully curated data sets where technical and biological confounders can be controlled for, is thus an important step in investigating the predictability of genetic signatures, as well as the identification of their morphological counterparts.

In most cases, such in-house datasets also contain technical and biological biases, due to the long period during which the dataset is acquired. This motivated us to propose a method to mitigate bias in Computational Pathology workflows, based on strategic sampling. Such strategies are already used in other fields of medical imaging but have so far - to the best of our knowledge - not been used in Computational Pathology. We have shown that this approach can successfully mitigate or even eliminate bias. However, the method is limited by the number of variables we can correct for, as well as by the class imbalance it can handle. In some cases, stratification might therefore be preferable. In any case, it is important to be aware of the confounding variables in the data set, whose presence can lead to false expectations and misinterpretation. For this reason, we expect proper treatment of such variables to become a standard in the field.

While bias correction on the TCGA led to a drop in AUC to 0.63, we found that HRD was predictable in our in-house data set of 715 BC patients with an AUC of 0.83. While homogeneous datasets do not reflect the variability between centers and thus limit direct applicability of the trained networks, they allow for controlled feasibility studies, which now need to be complemented by multicentric studies. In addition, we will validate this algorithm in a prospective neoadjuvant clinical trial for which patients’ HRD status will be assessed with MyChoice® CDx test (Myriad).

Homogeneous datasets are well suited for the identification of phenotypic patterns linked to disease patterns, even in cases where no such patterns are known a priori, such as in the case for HRD. In order to identify a phenotypic signature related to an output variable (here HRD), we can either use biologically meaningful encodings, also known as human interpretable features (HIF), and infer the most relevant features by analyzing the weights in the predictive model (8), or we can turn to network introspection. The HIF approach relies on detailed and exhaustive annotations of a large number of WSI. For instance, (8) leverage annotations provided by hundreds of pathologists consisting of hundreds of thousands of manual cell and tissue classifications. Here, we provide a new network introspection scheme, relying on the powerful MoCo encodings, trained without supervision directly on histopathology data, and a decision-based tile selection, that allows us to automatically cluster tiles and to relate these clusters to the output variable.

Interestingly, while our approach confirms the recently published finding that necrosis is a hallmark of HRD (8) and identifies morphological features common to HRD in TNBC and luminal BC, such as necrosis, high density in TILs and high nuclear anisokaryosis (48), it also points to more specific patterns that have so far been overlooked. For instance, we found tiles enriched in carcinomatous cells with clear cytoplasm suggesting activation of specific metabolic processes in these cells. Second, we find intra-tumoral laminated fibrosis as an HRD related pattern. This suggests the hypothesis that cancer-associated fibroblast (CAF) within the stroma of HRD luminal tumors may play a role in the viability and fate of tumor cells. Furthermore, the presence of adipose tissue within the tumor suggests first a different tumor cell density and second a specific balance between CAF and adipocytes in the context of a luminal HRD tumor. The molecular mechanisms achieving these patterns remain to be determined by in vitro models.

Similar to what we have shown here with respect to HRD, the visualization framework we have developed is versatile and can in principle be applied in the context of other genetic signatures. Because the algorithm is fully automated, using the MIL algorithm and its visualization method can constitute a useful tool for the discovery of morphological features related to the predicted genetic signatures. This has the potential to generate new biological hypotheses on the phenotypic impact of these genetic disorders. In order to maximize the benefit for the scientific community, we release the code to train MIL models on WSIs and to create morphological maps as well as tile trajectories publicly and free of charge, and provide detailed documentation.

Altogether, this study provides new and versatile tools for the prediction and phenotypic dissection of genetic signatures from histopathology data. Application to luminal breast cancers allowed us to shed light on the phenotypic consequences of homologous recombination deficiency, and might provide a tool with the potential to impact breast cancer patient care.

## Methods

### Data

#### In-house dataset (Institut Curie)

We retrospectively retrieved a series of 715 patients with HE slides of surgical resections specimens of untreated breast cancer and a genomically known HR status (Supplementary Table 1). The series is composed of 309 Homologous Recombination Proficient tumors (HRP) and 406 Homologous Recombination Deficient tumors (HRD). The HRD status was either identified by the presence of a germline *BRCA1/2* (*gBRCA1/2*) mutation or assessed by LST genomic signature according to Popova *et al*. for the sporadic triple-negative and luminal cancers.

All patients have been treated and followed at the Institut Curie between 1995 and 2020. The patient agreed for the use of tumor samples from their surgical resection specimens for research according to the law. Ethical approval from the Institutional Review Board (Institut Curie breast cancer study group N°DATA190031) was obtained for the use of all specimens. Clinical data have been retrieved from the Institut Curie electronic medical records and saved using Research electronic data capture (REDCap) tools hosted at the Institut Curie.

#### Public dataset (TCGA)

This public dataset is composed of 815 WSI of breast cancer fixed in formalin (FFPE) and stained in H&E. They are available at https://portal.gdc.cancer.gov/. Low-resolution WSI, WSI containing artifacts such as pen marks, tissue-folds and blurred WSI were removed. The final data-set encompasses 691 WSIs. The HR status of the corresponding tumors was obtained using the LST genomic signature (15).

#### Architecture and optimization parameters

Hyperparameters have been set thanks to a random search evaluated through 5-fold nested cross-validation. The benchmark task is the prediction of the molecular class of the TCGA WSIs. Both the decision module and the tile-scoring module are multilayer perceptrons with batch normalization (52) after each hidden layer. The decision module has 3 hidden layers of 512 neurons, the tile-scoring module has 1 hidden layer of 256 neurons.

Dropout has been fixed at 0.4, the optimizer is ADAM (53) with a learning rate of 3e-3. A batch consists of 16 samples of WSI. A sample of WSI corresponds to a uniform sampling of 300 of its composing tiles. In fact, we observed that this uniform subsampling of the WSIs regularized training as well as diminishes its computational workload. Finally, training is performed during 200 epochs.

Training and performance evaluation are done in a 5-fold nested cross-validation framework. Each dataset is split into 5 independent folds. For each of these folds, a validation set is randomly sampled in the complementary 4/5th. A model is trained on the remaining dataset (= ⅘ * ⅘ th of the total dataset). This process is repeated 10 times for each test fold, then the best model is selected according to its validation performances, and finally tested on its test set.

Each test and validation set preserves the stratification of the whole dataset with respect to the target variable as well as the confounding variables in case we correct for them. The final performance estimation of the model is the performance averaged over the 5 test performances. During inference time, all the tiles of each WSI are processed

#### Strategic sampling

is used both for balancing the training dataset with respect to the output variable (*T* ={*t*_1_,*t*_2_} in the binary case) and to correct for biases (*B* ={*b*_1_, *b*_2_, *b*_3_, .., *b*_*n*_}). If *X* is a given WSI sampled from the dataset, then *T(X)* and *B(X)* are respectively the target value and the bias value of *X*. We note |*t*_1_| the total number of slides in the dataset labeled with *t*_1_, same for |*b*_*i*_|. |*t*_*i*_*&b*_*i*_| is the total number of slides with label value *t*_*i*_ and bias value *b*_*i*_.

For achieving both, we sample the WSIs *X* in each batch in a distribution 𝒫 under which *𝒫*({*T*(*X*) = *t*_1_}) = *𝒫*({*T*(*X*) = *t*_2_}) and *𝒫*({*T*(*X*) = *t*_1_} ⋂ {*b*(*X*) = *b*_*i*_}) = *𝒫*({*T*(*X*) = *t*_2_} ⋂ {*b*(*X*) = *b*_*i*_}) for all i.

That is : *𝒫*(*X* | {*T*(*X*) = *t*_1_}⋂{*b*(*X*) = *b*_*i*_}) ∝ | *t*_1_ | / | *t*_1_ &*b*_*i*_ | for each *i* ⩽ *n, j* ⩽ *m*.

Strategic sampling is performed on the fly when building the batches.

When correction for several confounder simultaneously, *B* ={*b*_1_, *b*_2_, …, *b*_*n*_}, and *C* ={*c*_1_, *c*_2_, …, *c*_*m*_}, a new confounder variable is created that takes values in all the pairs combinations of *b*_*i*_ and 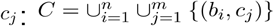. We then apply strategic sampling to correct for C.

#### Bias score

The bias score of a confounder variable *B* is the average mutual information between *B* and the predicted class *C* (I(*B*_*D*_,*C*_*D*_)) estimated in a dataset *D* in which the mutual information between *B* and the target *T* is zero.

We sample 30 sub-dataset *D*_*i*_ in the test set using strategic sampling such that 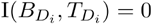 and average over them to get the bias score *BS*(*B*):

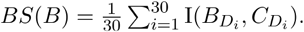

The mutual information *I*(*B,P*) between two finite random variables B and Q taking value respectively in {*b*_1_,,*b*_2_} and {*q*_1_, …, *q*_*m*_} measures the dependency between B and Q and is defined by :

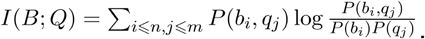

Because this average is non-negative even when *C* is not biased (but tends to zero when *D*_*i*_ is large), we compute the bias score of an unbiased model predicting the target variable with the same accuracy as the tested model. It serves as a control.

#### Learning MoCo representations

For learning MoCo-v2 representation we used the MoCo repository available at https://github.com/facebookresearch/moco.

We randomly used the following transformations: Gaussian blur, crop and resize, color jitter, grayscale, horizontal and vertical symmetries, and finally a color augmentation in the Hematoxylin and Eosin specific space (54).

The training dataset is composed of 5.3e6 images of size 224x 224 pixels, or half the Curie dataset at magnification 10x.

We used a Resnet18 and trained it for 60 epochs on 4 GPU Nvidia Tesla V100 SXM2 32 Go. We used the SGD optimizer with a momentum of 0.9, a weight decay of 1e-4 and a learning rate of 3e-3. We used a cosine scheduler with warm restart on the learning rate.

#### Visualization methods

The model used to extract the visualizations has been trained on the luminal subset of the Curie dataset (259 WSI). To benefit from the biggest dataset possible, the model has been trained on the whole dataset, without using early stopping nor testing, during 200 epochs. To generate the attention-based visualisation, the highest ranked tile with respect to the attention score is extracted, for each WSI. The selected tiles are then labeled according to the label of their WSI of origin.

Concerning the decision-based visualisation, for each WSI the 300 highest ranked tiles with respect to the attention score are selected. Amond this pool of tiles, the 2000 highest ranking tiles with respect to the posterior probability for HRD and HRP are selected. In order to promote diversity in the extracted images, no more than 20 tiles per slide can be selected.

#### Computation resources

All computations have been done on the GENCI HPC cluster of Jean-Zay.

## Code availability

The code that allows to reproduce this study and perform new ones is available at ******************************

## Supplementaries

Supplementaries are available at https://drive.google.com/file/d/1e1gEU8KfKl6LMq3lODWV3CJor9BgT_pE/view?usp=sharing

## Acknowledgements

The authors thank Hélène Guenon, Saida Sahiri Martial Caly and Laure Annette for their help in retrieving the H&E slides and their technical expertise. GB was supported by a Fondation Curie grant. TL was supported by a Q-Life PhD fellowship (Q-life ANR-17-CONV-0005). Furthermore, this work was supported by the French government under management of Agence Nationale de la Recherche as part of the ‘‘Investissements d’avenir’’ program, reference ANR-19-P3IA-0001 (PRAIRIE 3IA Institute).

## Authors contributions

AVS and GB initiated the project. AVS, GB, FCB and DSL generated the patient cohort. GB reviewed all the slides. TP and MHS performed the genomic analyses. TL, ED and TW designed the AI and statistical methods. TL and PN developed the software. TL performed the analysis, under the supervision of TW and ED. AVS and GB interpreted the morphological patterns. AVS, GB, TL, ED and TW discussed methods, results and design choices. TL prepared the figures. TL, TW and AVS wrote the manuscript.

## References

1. Veta M, Diest PJV, Willems SM, Wang H, Madabhushi A, Cruz-roa A, et al. Assessment of algorithms for mitosis detection in breast cancer histopathology images. Med Image Anal. 2014;1–23.

2. Ehteshami Bejnordi B, Veta M, Johannes van Diest P, van Ginneken B, Karssemeijer N, Litjens G, et al. Diagnostic Assessment of Deep Learning Algorithms for Detection of Lymph Node Metastases in Women With Breast Cancer. JAMA. 12 déc 2017;318(22):2199–210.

3. Campanella G, Hanna MG, Geneslaw L, Miraflor A, Silva VWK, Busam KJ, et al. Clinical-grade computational pathology using weakly supervised deep learning on whole slide images. Nat Med. août 2019;25(8):1301–9.

4. Mobadersany P, Yousefi S, Amgad M, Gutman DA, Barnholtz-Sloan JS, Velázquez Vega JE, et al. Predicting cancer outcomes from histology and genomics using convolutional networks. Proc Natl Acad Sci. 27 mars 2018;115(13):E2970–9.

5. Kather JN, Heij LR, Grabsch HI, Loeffler C, Echle A, Muti HS, et al. Pan-cancer image-based detection of clinically actionable genetic alterations. Nat Cancer. août 2020;1(8):789–99.

6. Coudray N, Ocampo PS, Sakellaropoulos T, Narula N, Snuderl M, Fenyö D, et al. Classification and mutation prediction from non–small cell lung cancer histopathology images using deep learning. Nat Med. oct 2018;24(10):1559–67.

7. Schmauch B, Romagnoni A, Pronier E, Saillard C, Maillé P, Calderaro J, et al. A deep learning model to predict RNA-Seq expression of tumours from whole slide images. Nat Commun [Internet]. 2020;11(1). Disponible sur: http://proxy.insermbiblio.inist.fr/login?url= http://search.ebscohost.com/login.aspx?direct=true&AuthType=ip,url,uid&db=edssjs&AN=edssjs.ADEC816B&lang=fr&site=eds-live

8. Diao JA, Wang JK, Chui WF, Mountain V, Gullapally SC, Srinivasan R, et al. Human-interpretable image features derived from densely mapped cancer pathology slides predict diverse molecular phenotypes. Nat Commun. déc 2021;12(1):1613.

9. Deluche E, Antoine A, Bachelot T, Lardy-Cleaud A, Dieras V, Brain E, et al. Contemporary outcomes of metastatic breast cancer among 22,000 women from the multicentre ESME cohort 2008–2016. Eur J Cancer. avr 2020;129:60–70.

10. Miller RE, Leary A, Scott CL, Serra V, Lord CJ, Bowtell D, et al. ESMO recommendations on predictive biomarker testing for homologous recombination deficiency and PARP inhibitor benefit in ovarian cancer. Ann Oncol Off J Eur Soc Med Oncol. déc 2020;31(12):1606–22.

11. Bryant HE, Schultz N, Thomas HD, Parker KM, Flower D, Lopez E, et al. Specific killing of BRCA2-deficient tumours with inhibitors of poly(ADP-ribose) polymerase. 2005;434:6.

12. Farmer H, McCabe N, Lord CJ, Tutt ANJ, Johnson DA, Richardson TB, et al. Targeting the DNA repair defect in BRCA mutant cells as a therapeutic strategy. Nature. avr 2005;434(7035):917–21.

13. Tung NM, Robson ME, Ventz S, Santa-Maria CA, Nanda R, Marcom PK, et al. TBCRC 048: Phase II Study of Olaparib for Metastatic Breast Cancer and Mutations in Homologous Recombination-Related Genes. J Clin Oncol. 29 oct 2020;38(36):4274–82.

14. Tutt ANJ, Garber JE, Kaufman B, Viale G, Fumagalli D, Rastogi P, et al. Adjuvant Olaparib for Patients with BRCA1-or BRCA2-Mutated Breast Cancer. N Engl J Med [Internet]. 3 juin 2021 [cité 17 juin 2021]; Disponible sur: http://www.nejm.org/doi/10.1056/NEJMoa2105215

15. Popova T, Manié E, Rieunier G, Caux-Moncoutier V, Tirapo C, Dubois T, et al. Ploidy and Large-Scale Genomic Instability Consistently Identify Basal-like Breast Carcinomas with BRCA1/2 Inactivation. Cancer Res. 1 nov 2012;72(21):5454–62.

16. Birkbak NJ, Wang ZC, Kim J-Y, Eklund AC, Li Q, Tian R, et al. Telomeric Allelic Imbalance Indicates Defective DNA Repair and Sensitivity to DNA-Damaging Agents. Cancer Discov. avr 2012;2(4):366–75.

17. Abkevich V, Timms KM, Hennessy BT, Potter J, Carey MS, Meyer LA, et al. Patterns of genomic loss of heterozygosity predict homologous recombination repair defects in epithelial ovarian cancer. Br J Cancer. nov 2012;107(10):1776–82.

18. Polak P, Kim J, Braunstein LZ, Karlic R, Haradhavala NJ, Tiao G, et al. A mutational signature reveals alterations underlying deficient homologous recombination repair in breast cancer. Nat Genet. oct 2017;49(10):1476–86.

19. Davies H, Glodzik D, Morganella S, Yates LR, Staaf J, Zou X, et al. HRDetect is a predictor of BRCA1 and BRCA2 deficiency based on mutational signatures. Nat Med. 1 avr 2017;23(4):517–25.

20. Tutt A, Tovey H, Cheang MCU, Kernaghan S, Kilburn L, Gazinska P, et al. Carboplatin in BRCA1/2-mutated and triple-negative breast cancer BRCAness subgroups: the TNT Trial. Nat Med. mai 2018;24(5):628–37.

21. Chopra N, Tovey H, Pearson A, Cutts R, Toms C, Proszek P, et al. Homologous recombination DNA repair deficiency and PARP inhibition activity in primary triple negative breast cancer. Nat Commun. déc 2020;11(1):2662.

22. Alexandrov LB, Nik-Zainal S, Wedge DC, Aparicio SAJR, Behjati S, Biankin AV, et al. Signatures of mutational processes in human cancer. Nature. août 2013;500(7463):415–21.

23. Chopra N, Tovey H, Pearson A, Cutts R, Toms C, Proszek P, et al. Homologous recombination DNA repair deficiency and PARP inhibition activity in primary triple negative breast cancer. Nat Commun. 29 mai 2020;11(1):2662.

24. Lakhani SR, Van De Vijver MJ, Jacquemier J, Anderson TJ, Osin PP, McGuffog L, et al. The pathology of familial breast cancer: predictive value of immunohistochemical markers estrogen receptor, progesterone receptor, HER-2, and p53 in patients with mutations in BRCA1 and BRCA2. J Clin Oncol Off J Am Soc Clin Oncol. 1 mai 2002;20(9):2310–8.

25. Manié E, Popova T, Battistella A, Tarabeux J, Caux-Moncoutier V, Golmard L, et al. Genomic hallmarks of homologous recombination deficiency in invasive breast carcinomas. Int J Cancer. 2016;138(4):891–900.

26. Ferrari A, Vincent-Salomon A, Pivot X, Sertier A-S, Thomas E, Tonon L, et al. A whole-genome sequence and transcriptome perspective on HER2-positive breast cancers. Nat Commun. 13 juill 2016;7(1):12222.

27. Turner NC. Signatures of DNA-Repair Deficiencies in Breast Cancer. N Engl J Med. 21 2017;377(25):2490–2.

28. Manié E, Popova T, Battistella A, Tarabeux J, Caux-Moncoutier V, Golmard L, et al. Genomic hallmarks of homologous recombination deficiency in invasive breast carcinomas: Genomic hallmarks of homologous recombination defect. Int J Cancer. 15 févr 2016;138(4):891–900.

29. Holstege H, Horlings HM, Velds A, Langerød A, Børresen-Dale A-L, van de Vijver MJ, et al. BRCA1-mutated and basal-like breast cancers have similar aCGH profiles and a high incidence of protein truncating TP53 mutations. BMC Cancer. déc 2010;10(1):654.

30. Ilse M, Tomczak JM, Welling M. Attention-based Deep Multiple Instance Learning. 180204712 Cs Stat [Internet]. 28 juin 2018 [cité 17 févr 2020]; Disponible sur: http://arxiv.org/abs/1802.04712

31. Amores J. Multiple instance classification: Review, taxonomy and comparative study. Artif Intell.1 août 2013;201:81–105.

32. Maron O, Lozano-Pérez T. A Framework for Multiple-Instance Learning. In: Jordan MI, Kearns MJ, Solla SA, éditeurs. Advances in Neural Information Processing Systems (NeurIPS). MIT Press; 1998. p. 570–6.

33. Courtiol P, Tramel EW, Sanselme M, Wainrib G. Classification and disease localization in histopathology using only global labels: a weakly supervised approach. CoRR. 2017;1–13.

34. He K, Fan H, Wu Y, Xie S, Girshick R. Momentum Contrast for Unsupervised Visual Representation Learning. 191105722 Cs [Internet]. 23 mars 2020 [cité 17 mars 2021]; Disponible sur: http://arxiv.org/abs/1911.05722

35. Valieris R, Amaro L, Osório CAB de T, Bueno AP, Rosales Mitrowsky RA, Carraro DM, et al. Deep Learning Predicts Underlying Features on Pathology Images with Therapeutic Relevance for Breast and Gastric Cancer. Cancers. déc 2020;12(12):3687.

36. Kather JN, Pearson AT, Halama N, Jäger D, Krause J, Loosen SH, et al. Deep learning can predict microsatellite instability directly from histology in gastrointestinal cancer. Nat Med. juill 2019;25(7):1054–6.

37. Kleppe A, Skrede O-J, De Raedt S, Liestøl K, Kerr DJ, Danielsen HE. Designing deep learning studies in cancer diagnostics. Nat Rev Cancer. 29 janv 2021;1–13.

38. Varoquaux G, Raamana PR, Engemann D, Hoyos-Idrobo A, Schwartz Y, Thirion B. Assessing and tuning brain decoders: cross-validation, caveats, and guidelines. NeuroImage. janv 2017;145:166–79.

39. Zhao Q, Adeli E, Pohl KM. Training confounder-free deep learning models for medical applications. Nat Commun. 26 nov 2020;11(1):6010.

40. Zhao J, Wang T, Yatskar M, Ordonez V, Chang K-W. Men Also Like Shopping: Reducing Gender Bias Amplification using Corpus-level Constraints. 170709457 Cs Stat [Internet]. 28 juill 2017 [cité 23 févr 2021]; Disponible sur: http://arxiv.org/abs/1707.09457

41. Adeli E, Zhao Q, Pfefferbaum A, Sullivan EV, Fei-Fei L, Niebles JC, et al. Representation Learning with Statistical Independence to Mitigate Bias. 191003676 Cs [Internet]. 20 nov 2020 [cité 3 févr 2021]; Disponible sur: http://arxiv.org/abs/1910.03676

42. Wang T, Zhao J, Yatskar M, Chang K-W, Ordonez V. Balanced Datasets Are Not Enough: Estimating and Mitigating Gender Bias in Deep Image Representations. 181108489 Cs [Internet]. 10 oct 2019 [cité 8 févr 2021]; Disponible sur: http://arxiv.org/abs/1811.08489

43. Wang Z, Qinami K, Karakozis IC, Genova K, Nair P, Hata K, et al. Towards Fairness in Visual Recognition: Effective Strategies for Bias Mitigation. In: 2020 IEEE/CVF Conference on Computer Vision and Pattern Recognition (CVPR) [Internet]. Seattle, WA, USA: IEEE; 2020 [cité 18 févr 2021]. p. 8916-25. Disponible sur: https://ieeexplore.ieee.org/document/9156668/

44. Lu MY, Williamson DFK, Chen TY, Chen RJ, Barbieri M, Mahmood F. Data-efficient and weakly supervised computational pathology on whole-slide images. Nat Biomed Eng. 1 mars 2021;1–16.

45. Dehaene O, Camara A, Moindrot O, de Lavergne A, Courtiol P. Self-Supervision Closes the Gap Between Weak and Strong Supervision in Histology. 201203583 Cs Eess [Internet]. 7 déc 2020 [cité 31 mars 2021]; Disponible sur: http://arxiv.org/abs/2012.03583

46. Courtiol P, Maussion C, Moarii M, Pronier E, Pilcer S, Sefta M, et al. Deep learning-based classification of mesothelioma improves prediction of patient outcome. Nat Med. oct 2019;25(10):1519–25.

47. McInnes L, Healy J, Melville J. UMAP: Uniform Manifold Approximation and Projection for Dimension Reduction. 180203426 Cs Stat [Internet]. 6 déc 2018 [cité 19 nov 2019]; Disponible sur: http://arxiv.org/abs/1802.03426

48. Rakha EA, El-Sayed ME, Reis-Filho J, Ellis IO. Patho-biological aspects of basal-like breast cancer. Breast Cancer Res Treat. 1 févr 2009;113(3):411–22.

49. Stratton MR. Pathology of familial breast cancer: differences between breast cancers in carriers of BRCA1 or BRCA2 mutations and sporadic cases. The Lancet. 24 mai 1997;349(9064):1505–10.

50. Lakhani SR, Gusterson BA, Jacquemier J, Sloane JP, Anderson TJ, van de Vijver MJ, et al. The Pathology of Familial Breast Cancer: Histological Features of Cancers in Families Not Attributable to Mutations in BRCA1 or BRCA2. Clin Cancer Res. 1 mars 2000;6(3):782.

51. Bane AL, Beck JC, Bleiweiss I, Buys SS, Catalano E, Daly MB, et al. BRCA2 Mutation-associated Breast Cancers Exhibit a Distinguishing Phenotype Based on Morphology and Molecular Profiles From Tissue Microarrays. Am J Surg Pathol [Internet]. 2007;31(1). Disponible sur: https://journals.lww.com/ajsp/Fulltext/2007/01000/BRCA2_Mutation_associated_Breast_Cancers_Exhibit_a.15.aspx

52. Ioffe S, Szegedy C. Batch Normalization: Accelerating Deep Network Training by Reducing Internal Covariate Shift. 150203167 Cs [Internet]. 10 févr 2015 [cité 20 oct 2019]; Disponible sur: http://arxiv.org/abs/1502.03167

53. Kingma DP, Ba J. ADAM: A Method for Stochastic Optimization. 14126980 Cs [Internet]. 29 janv 2017 [cité 21 avr 2021]; Disponible sur: http://arxiv.org/abs/1412.6980

54. Ruifrok AC. Quantification of histochemical staining by color deconvolution. :21.

